# Risk to North American Birds from Climate Change-Related Threats

**DOI:** 10.1101/798694

**Authors:** Brooke L. Bateman, Lotem Taylor, Chad Wilsey, Joanna Wu, Geoffrey S. LeBaron, Gary Langham

## Abstract

Climate change is a significant threat to biodiversity globally, compounded by threats that could hinder species’ ability to respond through range shifts. However, little research has examined how future bird ranges may coincide with multiple stressors at a broad scale. Here, we assess the risk to 544 birds in the United States from future climate change threats under a mitigation-dependent global warming scenario of 1.5°C and an unmitigated scenario of 3.0°C. Threats considered included sea level rise, lake level change, human land cover conversion, and extreme weather events. We developed a gridded index of risk based on coincident threats, species richness, and richness of vulnerable species. To assign risk to individual species and habitat groups, we overlaid future bird ranges with threats to calculate the proportion of species’ ranges affected in both the breeding and non-breeding seasons. Nearly all species will face at least one new climate-related threat in each season and scenario analyzed. Even with lower species richness, the 3.0°C scenario had higher risk for species and groups in both seasons. With unmitigated climate change, multiple coincident threats will affect over 88% of the conterminous United States, and 97% of species could be affected by two or more climate-related threats. Some habitat groups will see up to 96% species facing three or more threats. However, climate change mitigation would reduce risk to birds from climate change-related threats across over 90% of the US. Across the threats included here, extreme weather events have the most significant influence on risk and the most extensive spatial coverage. Urbanization and sea level rise will also have disproportionate impacts on species relative to the area they cover. By incorporating threats into predictions of climate change impacts, this assessment provides a comprehensive picture of how climate change will affect birds and the places they need.

## INTRODUCTION

Climate change is a significant threat to biodiversity globally, and species must find ways to cope with changing environmental conditions [1,2]. Range shifts are expected to be a crucial response to a changing climate, with some species tracking suitable environmental conditions as they move across the landscape [3–5]. With the rapid pace of contemporary climate change, many species will need to shift faster and farther to keep pace [6,7]. Predictions of range shifts are typically conducted with species distribution models (SDM), which pair species occurrence data with projected climate and environmental information to map species’ ranges under climate change [8,9]. However, range shifts are only part of the climate change story: multiple coincident threats, like extreme weather events and land-use change, are another factor that compounds the risk of climate change [10], since species may face additional obstacles as they move across the landscape in search of suitable conditions [3], which could hinder their ability to respond to climate change through range shifts. Despite this, SDM-based prioritizations rarely consider additional threats that put biodiversity at risk [11]. Focusing solely on climate change and not including other threats could severely underestimate extinction risk in the future [3,12,13].

Here, we assess the risk of birds in the conterminous United States (US) to multiple climate-related threats, incorporating potential future range shifts. Birds as a group are particularly sensitive to climate change [1,3,14]. Indeed, there is much evidence that birds have already responded to contemporary climate change through range shifts [15–20]. However, there has been little research on how future projections of bird ranges may coincide with multiple stressors at a broad scale. Bird SDM outputs based on climate are sometimes assessed along with single threats like land-use change [1,12,21,22], but rarely is more than one threat considered.

Climate-related threats can arise directly from changes to a climate regime, such as extreme weather events like extreme heat, droughts, fire weather, false springs, and heavy rainfall, where conditions range outside the norm in terms of magnitude and frequency. There is evidence that these intermittent events will become stronger and more frequent with climate change [23,24]. Birds may be especially vulnerable to the haphazard and abrupt nature of extreme events, which can drastically reduce population numbers [25]. Extreme weather can have significant effects on populations through direct impacts on vital rates and indirect impacts on habitat selection and resource availability [26,27]. For example, extreme heat can directly affect individuals through heat stress [27], which can lead to mass mortality events [28], and subsequently reduce populations and species richness locally [29]. Similarly, droughts can also cause mortality [27,30], diminish reproductive success [31], lead to declines in abundance and richness [32], and trigger species movement [33]. At the other end of the precipitation spectrum, heavy rainfall can flood nests and burrows, killing chicks [27,34–38]. False springs are a less apparent and acute threat, and occur when a hard freeze follows premature warm temperatures in late winter or early spring. They can cause vegetation damage and cascading ecosystem effects [39,40], including reduced food sources for primary consumers [41], which then limits the resources available to feed young [42]. In isolation, extreme weather events are well-documented to have negative consequences. However, a given species will likely face multiple events locally [23,26,43,44]. The combined influence of these events can dramatically alter populations and communities [26,45], and cause physiological stress to birds [26].

Threats such as fire, sea level rise, and lake level change may also be catastrophic, causing a change in ecosystem state that radically alters habitat quality and availability. Fire is a complex process that, while sometimes destructive in its immediate effects, can also play an essential role in maintaining habitat over time. The impacts of fire vary depending on multiple factors, including species response, habitat type, and severity, but are generally damaging in the short-term, leading to mortality, displacement, and population declines for some species [46]. Birds may return to burned areas as vegetation regenerates and provides diverse foraging and nesting resources for many species [47].

In contrast, sea level rise is likely to cause catastrophic habitat loss over the long-term, as nesting sites become inundated or transition to different habitat types [48,49]. Coastal species will also suffer in the short-term, as flooding becomes more frequent and catastrophic from storm tides [23,25], leading to direct mortality of chicks as nests and burrows are destroyed [25]. The unpredictable pattern of these changes may also create ecological traps, as birds are unable to adapt or move to new areas [25]. Similarly, changes in lake level variability, primarily associated with reductions in lake levels, can alter shoreline and wetland habitats and their ecological function [50–52]. Frequently, waterbird and shorebird species rely upon specific wetland vegetation communities, and changes in seasonal hydrology can reduce reproductive success through fewer breeding pairs and more nest failures [53].

Some threats may also be indirectly related to climate change, such as changes in human land use, including cropland expansion and urbanization. These changes have already created a fragmented and degraded landscape, with estimates of more than 50% of the earth’s land surface modified for human use [54] and intensification of human land-use change likely in the future [55,56]. In the conterminous US, we have already seen rapid rates of land-use conversion in both agricultural and urban expansion with human population growth. Loss and degradation of habitats due to human activity and land-use change are a significant threat to biodiversity, including birds [1,21,57]. Bird species richness and abundance are both negatively associated with anthropogenic land use [58–60], especially for habitat specialists and species of conservation concern [61]. Areas with increased pressure from both land-use change and climate change have also seen accelerated losses in bird populations and communities [12]. The combined and additive effects of land use and climate change will likely lead to high rates of biodiversity loss, a homogenization of communities, and reduced ecosystem functioning [62,63].

The goal of our analysis was to identify the places and birds most at risk in the conterminous United States under two global warming scenarios, 1.5°C and 3.0°C. We chose global warming scenarios of 1.5°C and 3.0°C to represent a range of climate futures that are almost certain to occur. The most recent IPCC report identifies 1.5°C global mean temperature rise to be nearly inevitable within the next few decades without swift and aggressive policy changes to reduce greenhouse gases [24], and the Paris Agreement includes pledged reductions in greenhouse gas emissions leading to an estimated 3.2°C global increase in mean temperature [64]

## MATERIALS AND METHODS

### Framework

We estimated the local risk to birds of climate-related threats based on the IUCN framework described by Foden & Young (2016). The IUCN defines risk as the probability of harmful consequences resulting from climate change based on the interaction of (1) hazard, the potential occurrence of a damaging physical event, (2) exposure, the presence of species or systems that could be adversely affected, and (3) vulnerability, the predisposition to be adversely affected based on sensitivity and lack of adaptive capacity [65]. In our assessment, hazard was based on future threats, including sea level rise, lake level change, land cover conversion from natural to human use, and five types of extreme weather events: extreme spring heat, fire weather, spring droughts, heavy precipitation, and false springs. Exposure and vulnerability were based on a previous analysis modeling North American bird species’ ranges and assessing their climate change vulnerability [66]. We assessed places at risk by developing a gridded index of risk as the product of hazard, exposure, and vulnerability (Fig 1), and we assessed species at risk by calculating the proportion of each species’ future range that overlaps with each climate-related threat.

**Fig 1.**
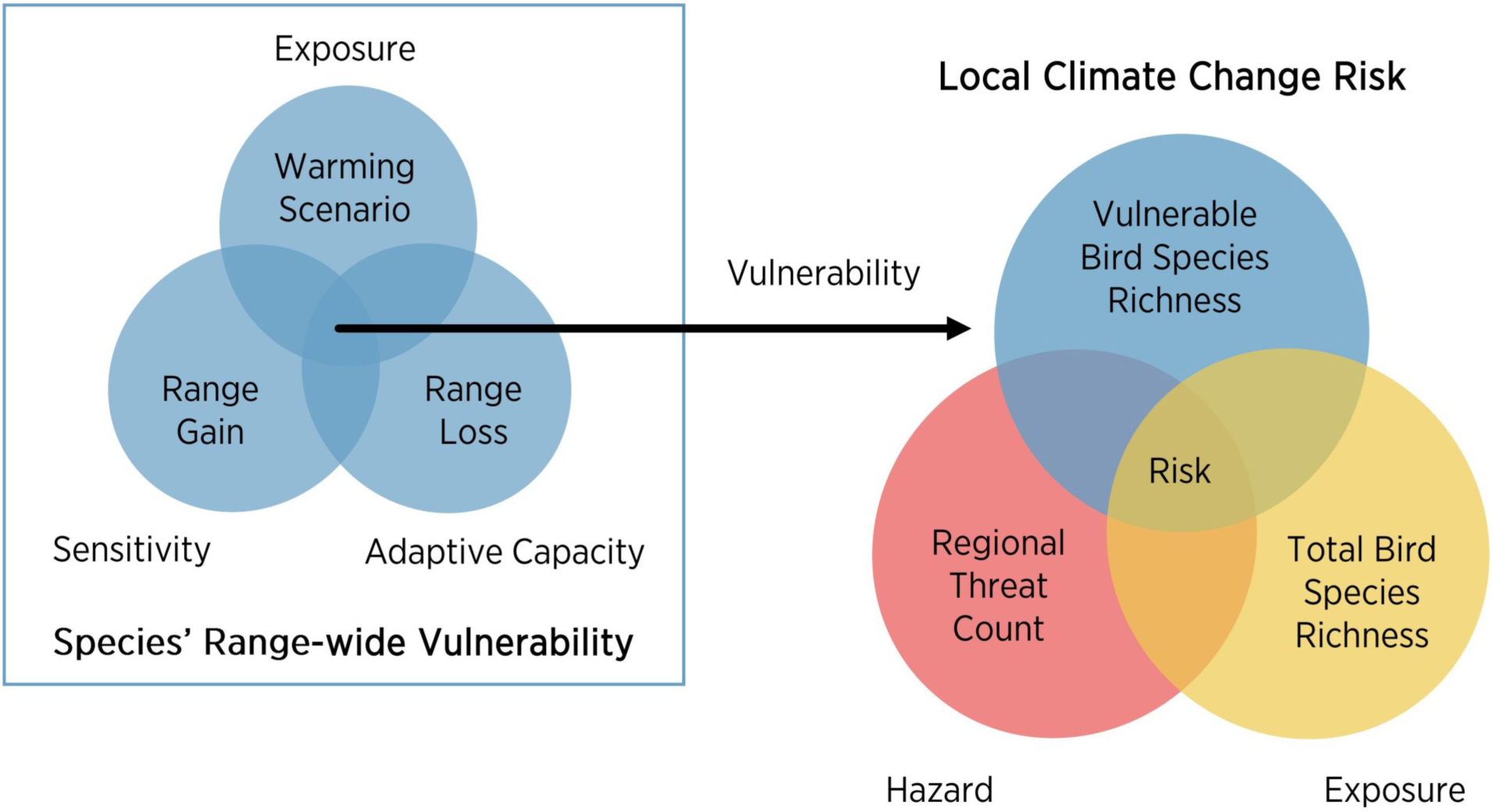
Risk assessment framework. Species vulnerability is a function of exposure to change (measured here by global warming scenario), sensitivity to change (range loss), and adaptive capacity in the face of change (range gain relative to range loss). Risk is a function of exposure (number of species), vulnerability (number of vulnerable species), and hazard (coincident threats). Adapted from Foden and Young [65].

### Data

We assembled gridded datasets for the conterminous US of climate change-related threats, along with projections of bird species’ ranges under both a 1.5°C and 3.0°C mean global temperature rise scenario. We equated projections from representative concentration pathway (RCP) 4.5 at mid-century (2041-2070, or 2050 for decadal outputs) with the 1.5°C global mean temperature rise scenario and RCP8.5 at end of century (2071-2100, or 2100 for decadal outputs) with a 3.0°C global mean temperature rise scenario [67]. RCP4.5 assumes global greenhouse gas emissions will level off and stabilize by end of century, while RCP8.5 assumes that emissions will continue to increase through 2100 [68]. We aligned all datasets to a common gridded format covering the conterminous US at a 1-km resolution, using nearest neighbor resampling for coarser resolution datasets (i.e. > 1-km) and average resampling for finer resolution datasets (i.e. < 1-km) in order to calculate the proportion of each 1-km cell affected by each threat. All projections of the data were based on an ensemble general circulation model (GCM) when available, or averaged across available GCMs otherwise.

### Sea Level Rise

We mapped areas of sea level r ise-induced flooding and ensuing habitat transitions based on spatial projections of sea level rise and associated marsh migration from NOAA’s Office for Coastal Management (available from https://coast.noaa.gov/slr/). Projections are based on a modified bathtub approach that incorporates LIDAR-derived elevation data and attempts to account for local and regional tidal variability [49]. Outputs are available for the conterminous US at a 10-m spatial resolution with scenarios of up to 10 feet (∼3 m) provided in half-foot increments. We selected scenarios of 0.5 m and 1.0 m, which crosswalk with estimates from the IPCC for 1.5°C and 3.0°C warming scenarios by end of century [69], as well as an additional “extreme” scenario (2.0 m) to capture current high-end estimates of sea level rise under 3.0°C [70–72]. We then downscaled these global scenarios to states using a dataset from the National Climate Assessment that localizes global sea level rise projections [73]. Downscaled estimates are available for six sea level rise scenarios (0.3, 0.5, 1.0, 1.5, 2.0, and 2.5 m) at 263 locations within the 22 coastal states in the conterminous US (mean=12 locations per state, range=2-44). Because downscaling by state resulted in variations of over 1.0 m (i.e. the 2.5 m scenario could results in state-level SLR over 3.5m), the 2.0 m scenario was the highest scenario we could include to maintain our ability to match downscaled estimates to spatial projections (which were only available for scenarios of up to 10 feet, ∼3 m).

### Lake Level Change

We obtained spatial projections of lake level change for the Laurentian Great Lakes from NOAA’s Office for Coastal Management (available from https://coast.noaa.gov/llv/). Estimates of lake extent within the US are available for scenarios of -6 to +6 feet of change based on LI-DAR-derived topographic and bathymetric elevation data. Lake levels are generally expected to decline under climate change as (1) surface water temperatures warm, increasing rates of evaporation, and (2) lake ice forms later, extending the season for evaporation [74]. However, lake levels also vary considerably [74]. Therefore, we summed historical low and high water levels with projections of future mean water levels for each lake to identify areas that could be affected by both drying (with low lake levels) and flooding (with high lake levels) in the future. Because minimums and maximums can be subject to outliers, we calculated the 1st and 99th percentile of water levels between 1860 and 2015 to estimate historical low and high water levels for each lake based on shoreline gauging data from NOAA’s Great Lakes Environmental Research Laboratory (available from www.glerl.noaa.gov). Lake St. Clair was also included, and lakes Michigan and Huron were grouped because they are connected at the same water level. We summed historical variability with estimated mean water levels for Michigan-Huron for our two global warming scenarios from a previous assessment [75]. Estimates of long-term change for the other lakes are negligible under the assumption that water regulation practices will continue [75]. We used our summed estimates to select a low and high water level projection for each lake, and then combined these spatial projections (removing the current lake extent) to identify the total area prone to drying or flooding under our two climate change scenarios.

### Human Land Use Change

We considered two types of land cover conversion from natural to human use: urbanization and cropland expansion. We obtained projections of urban growth from the EPA’s Integrated Climate and Land Use Scenarios (ICLUS) dataset (available from https://iclus.epa.gov). ICLUS projections are derived from a pair of models: a demographic model generates county-level population estimates, and a spatial allocation model distributes new urban development in response to population growth. Outputs are available for the conterminous US at a 90-m spatial resolution for every decade between 2000 and 2100 under two general circulation models (GISS-E2-R, HadGEM2-ES) and two climate change scenarios combining shared socioeconomic pathways (SSPs) and RCPs (SSP2+RCP4.5 and SSP5+RCP8.5). We equated these scenarios at 2050 and 2100, respectively, to our two global warming scenarios, 1.5° and 3.0°C. Projections of cropland expansion were obtained from a downscaled version of the Land-Use Harmonization (LUH) dataset [76], a 0.5° resolution gridded dataset spanning the years 1500–2100 that estimates urban and agricultural land use patterns and transitions [77]. A previous assessment downscaled this dataset to a 1-km resolution using a Cellular Automata approach, and outputs were generated for every decade between 2010 and 2100 under four representative concentration pathways [RCP2.6, RCP4.5, RCP6.0, and RCP8.5;, 76]. We utilized projections from RCPs 4.5 and 8.5 at 2050 and 2100, respectively, for our 1.5° and 3.0°C scenarios.

### Extreme Weather

We considered five extreme weather variables: extreme spring heat, spring droughts, fire weather, heavy rain, and false springs. Spatial projections were obtained from previous assessments [39,43,44; available from http://silvis.forest.wisc.edu/climate-averages-and-extremes/] that derived these variables from daily projections of the 19 GCMs participating in the Coupled Model Intercomparison Project 5 (CMIP5), statistically downscaled to a 12-km resolution using the Bias-Corrected Constructed Analog technique [78]. Again we used projections from two RCPs (4.5 and 8.5) aligned with mid- and late-century, respectively for our climate change scenarios. Extreme spring heat and spring droughts were calculated based on standardized indices, the standard temperature index [STI; 43] and precipitation index [SPI; 79], respectively, that define extreme events based on a standard normal distribution, with outputs representing the frequency of 20-year extreme events, and reciprocal values (i.e. 1/x) representing the return interval [43]. Fire weather [based on the Keetch-Byram Drought Index, KBDI;, 80] and heavy rain were calculated as the number of days above the 95th percentile of historical values [44]. Fire weather, a measure of soil moisture deficit and drought indicative of wildfire potential, provides a reference for weather conditions suitable for fire in a given season, but does not directly translate into more fires because fire events require not only appropriate weather, but also fuels, ignition sources, an topography [81]. False springs were calculated as the probability of a hard freeze after leaf and flower emergence [39]. We mapped areas affected by these extreme weather events by applying thresholds to convert continuous values into categorical outputs. We explored multiple thresholds of more frequent return intervals than 20-years for extreme spring heat and spring droughts, increased number of days above the historical values for fire weather and heavy rain, and increase in the probability of false springs. From this, we mapped the following areas: extreme spring heat happening every 2 years or more often in the future, spring droughts happening every 10 years or more often in the future, 90 days of the year or more exceeding the current 95th percentile of fire weather, 10 days of the year or more exceeding the 95th percentile of precipitation, and a 50% or greater chance of a false spring in the future.

### Birds

We incorporated predictions of future bird ranges in the conterminous US to characterize bird exposure and vulnerability to climate change in our risk assessment. Bateman et al. [66] developed species distribution models for 604 species (including 597 breeding and 545 non-breeding season models) by relating bird observations to a set of bioclimatic and habitat variables. These models were based on the methodology of Wilsey et al for 38 grassland birds, applied to an additional 566 species in North America [67]. Each species was assigned to one of 12 habitat groups (arctic, aridlands, boreal forests, coastal, eastern forests, generalists, grasslands, marshlands, subtropical forests, urban/suburban, waterbirds, and western forests), and modeled in a group-based approach. The complete bird observation dataset included more than 140 million georeferenced records from nearly 70 datasets. Bioclimatic and habitat variables for all species in both seasons included climatic moisture deficit, number of frost-free days, mean annual precipitation, precipitation as snow, vegetation type, terrain ruggedness, and anthropogenic land cover. Additional bioclimatic variables for the breeding season included mean temperature of the warmest month, chilling degree days, and summer heat moisture index. Additional bioclimatic variables for the nonbreeding season included mean temperature of the coldest month and growing degree days. Group-specific habitat variables included surface water (marshlands and waterbirds); wetland type (marshlands and waterbirds); distance to wetlands (waterbirds); distance to coast, excluding inland water bodies (coastal); distance to shore, including inland water bodies (marshlands and waterbirds); and a human influence index (urban/suburban).

Continuous model projections were converted to binary range maps by applying a threshold selected from multiple statistical metrics including mean occurrence prediction (mo; mean suitability prediction for the occurrence records), maximum sensitivity specificity (tss; maximized sensitivity + specificity), 10% omission (om_10; excludes 10% of occurrence records), sensitivity specificity (eq; sensitivity is equal to specificity), maximum Kappa (mk; maximum Kappa statistic), minimum occurrence prediction, (min_pred; minimum suitability prediction across all occurrence records), and a derived custom threshold to fall between the minimum occurrence prediction and 10% omission thresholds ((om_10+ min_pred)/3). Final threshold was selected based on threshold statistics and then reviewed via expert opinion, whereas experts verified each species/season model combination approximated ecological reality for that species. While species distribution models covered the full continental US, Canada, and Mexico, the scope of this assessment was limited to the conterminous US, resulting in truncated species ranges at political borders. We included species with at least 5% of their modeled range in the conterminous US, resulting in 544 species considered in this assessment, including 450 breeding (131 vulnerable) and 467 (74 vulnerable) non-breeding species present under a 1.5°C warming scenario, and 409 breeding (165 vulnerable) and 479 non-breeding (118 vulnerable) species present under a 3.0°C warming scenario.

Bateman et al. [66] also assessed vulnerability for each species within each season and scenario based on projections of range loss and potential range gain using the framework from Wilsey et al. [67]. This framework uses the proportion of projected loss from the current range as a measure of sensitivity, and the ratio of projected current range loss to projected future range gain as a measure of adaptive capacity. These two measures were summed for a final vulnerability score, classified into neutral, low, moderate, or high. We considered species in the moderate and high vulnerability classes to be vulnerable to climate change.

### Risk Assessment

Here, risk is defined as the product of hazard, exposure, and vulnerability [65]. For each season and scenario, we developed a gridded index of risk by first calculating each component of risk and then multiplying them together (Fig 1). To make results more directly comparable, we rescaled risk from 0-1 based on the minimum and maximum values across seasons and scenarios.

Hazard is defined as a climate-related event or trend that has potential to damage an ecosystem [65]. We measured hazard as the coincident number of threats across the conterminous US, calculated as the sum of each of our nine climate-related threats at both a 1.5°C and 3.0°C global warming scenario, and assessed the area affected by multiple threats. For threats with an original resolution finer than 1-km (i.e. sea level rise, lake level change, urbanization), we included all 1-km cells with at least 25% coverage of that 1-km cell.

To measure exposure, defined as the presence of species that could be adversely affected by a threat [65] we used modeled future range maps to estimate richness for the breeding and non-breeding seasons at 1.5°C and 3.0°C [66].

To measure vulnerability, defined as species that are prone to be adversely affected based on sensitivity and lack of adaptive capacity [65], we used modeled future range maps to estimate richness of only highly or moderately vulnerable species [66]. Because the risk index is a product, we offset vulnerability values by +1 so that risk in areas with no vulnerable species would be based on hazard and exposure only.

To summarize impacts on individual species, we calculated the area of each species’ projected range that coincides with each threat, relative to the species’ total conterminous US range. For threats that will result in long-term persistent changes to bird habitat (i.e. sea level rise, lake level change, human land use change), we considered species affected if at least 10% of their range overlapped with the threat. For threats that will result in short-term or intermittent changes to habitat (i.e. extreme weather events, including fire weather), we considered species affected if at least 50% of their range overlapped with the threat. These thresholds were selected after a sensitivity analysis across all species ranges and overlapping threats, and full details for each species, season, scenario, and threat along with the amount of area that each threat covers in a species range and the proportion of that threat will be provided in the supplementary information. For each season and scenario, we summed these results to identify the total number of species affected by each threat, and the number of vulnerable species affected. We then summed the number of threats affecting each species to identify the number and proportion of species that were affected by multiple coincident threats. We also report species results broken down by the proportion of species affected by each threat within our bird habitat groups.

## RESULTS

### Threats: Hazard

Overall, short-term intermittent threats (i.e. extreme weather) covered a much greater area than long-term persistent threats (i.e. sea level rise, lake level change, and human land use change) (Fig 2, see S1 Fig for maps of single threats). Extreme spring heat was the most ubiquitous threat, covering over 98% of the conterminous US under 3.0°C of warming and over half of this area (51%) under 1.5°C of warming; whereas sea level rise and lake level change were the most geographically restricted threats, covering 1% or less of the conterminous US under both scenarios (Fig 2). Fire weather was another substantial threat under 3.0°C, affecting about two-thirds of the conterminous US. Nearly all threats covered more area under 3.0°C compared to 1.5°C; the one exception was cropland expansion, which affected a small area under both scenarios, but covered nearly three times more area under 1.5°C compared to 3.0°C (2.3% vs 0.8% of the conterminous US; Fig 2).

**Fig 2.**
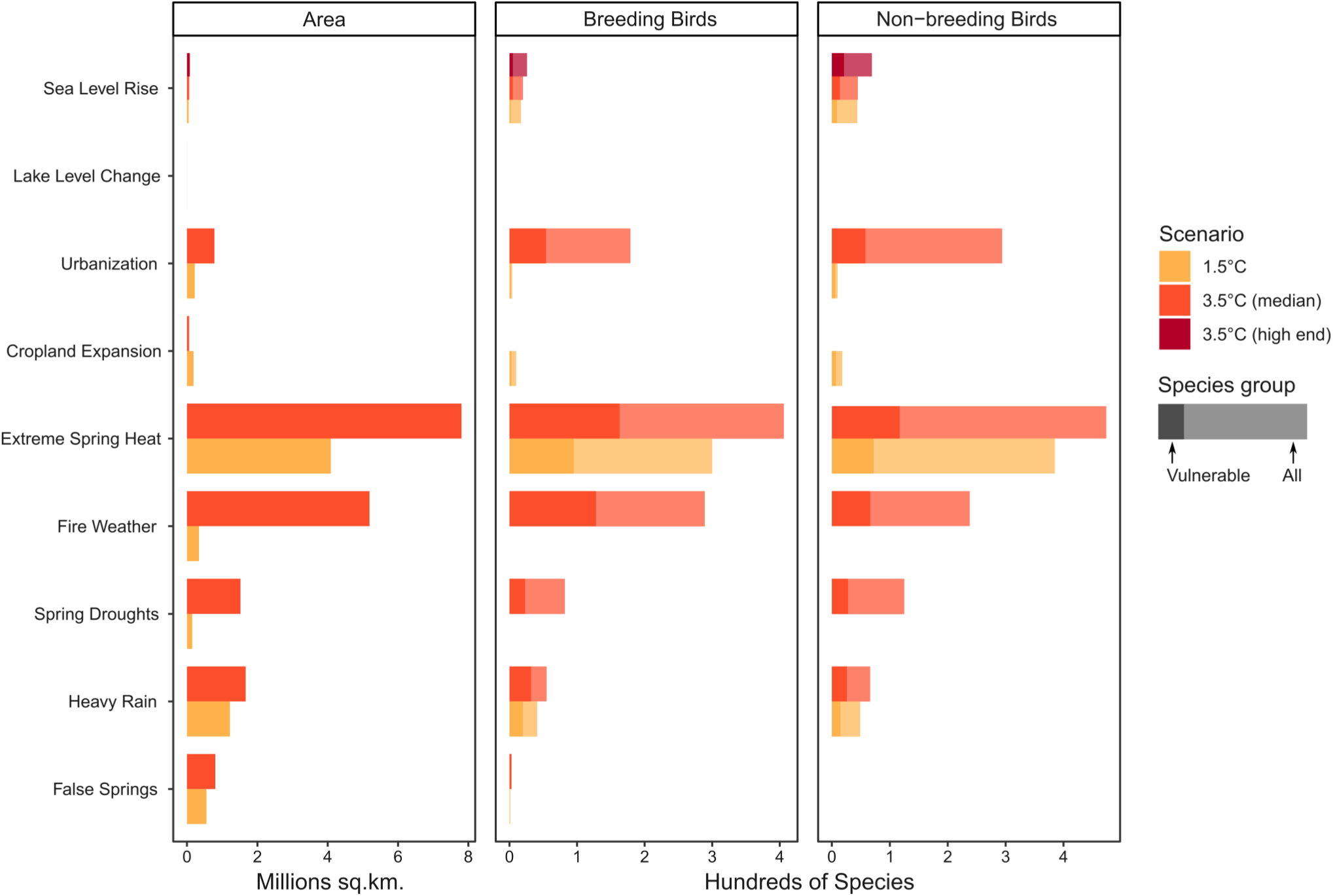
Area and number of species affected by threats under future global change scenarios of 1.5°C and 3.0°C (including a median and high-end option for sea level rise). Affected species overlap with threats in at least 10% (for persistent threats) or 50% (for intermittent threats) of their range in the conterminous US.

Regardless of season, climate-related threats are more widespread and intense under a 3.0°C scenario. Under a 3.0°C scenario, some locations might experience up to six of the nine threats included in this analysis, with over 98% of the US affected by at least one threat, and over 88% affected by two or more threats (Figs 3-4). Threats were most concentrated in the Northeast (from West Virginia north to Maine), Southwest (from Texas west to Arizona), and Gulf Coast (Louisiana) and least concentrated in the southern Interior Lowlands and Coastal Plains. Climate change mitigation to reduce warming from a 3.0°C to 1.5°C would slightly reduce the maximum number of coincident threats from 6 to 5 threats/km2, but the area projected to experience climate-related threats would be substantially reduced (Fig 4). Under a 1.5°C scenario, over 66% of the conterminous US would experience at least one threat, and only 19% would be affected by two or more threats (Fig 3).

**Fig 3.**
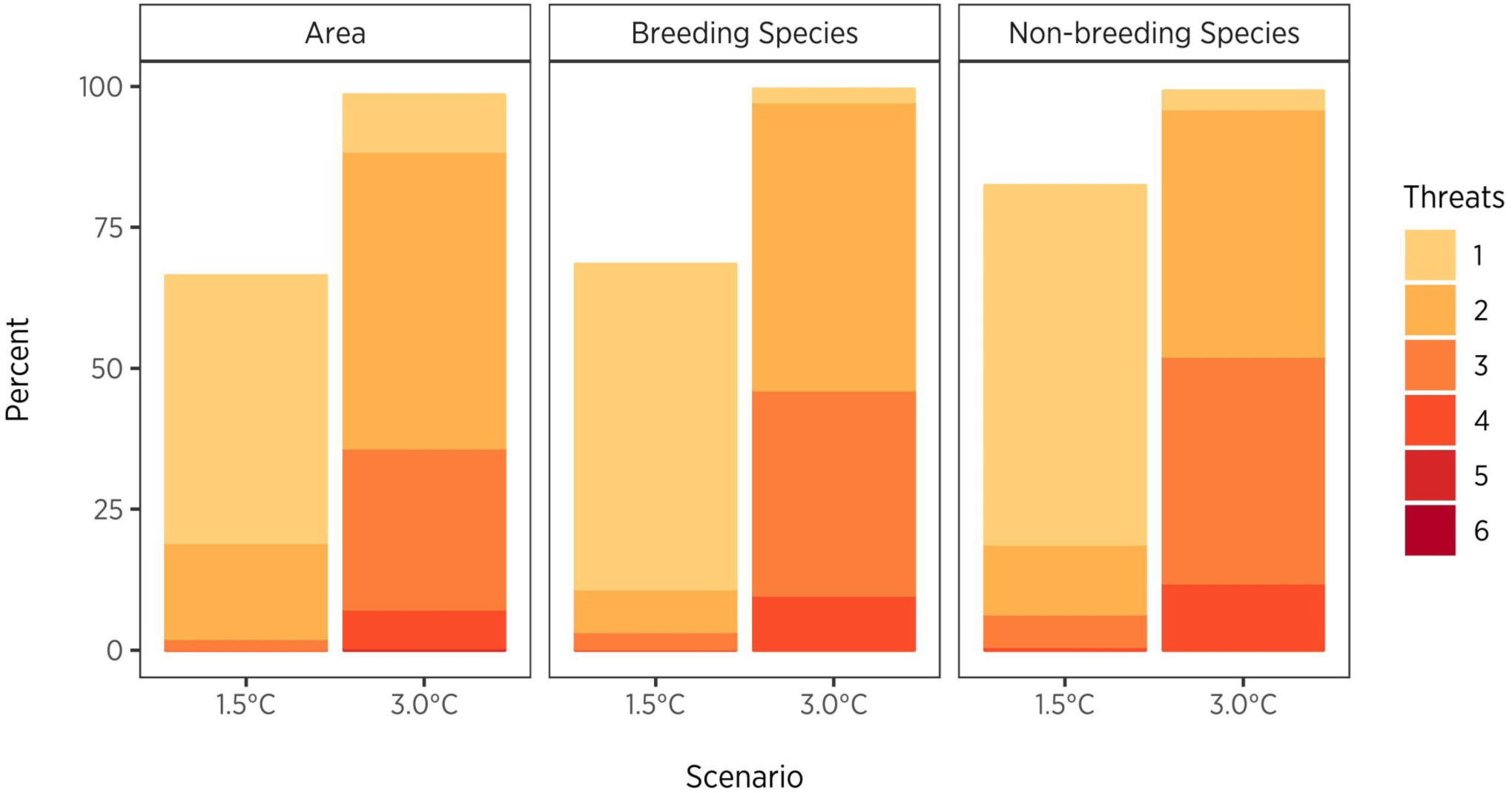
Proportion of area and breeding and non-breeding species affected by coincident number of threats under future global change scenarios of 1.5°C and 3.0°C.

**Fig 4.**
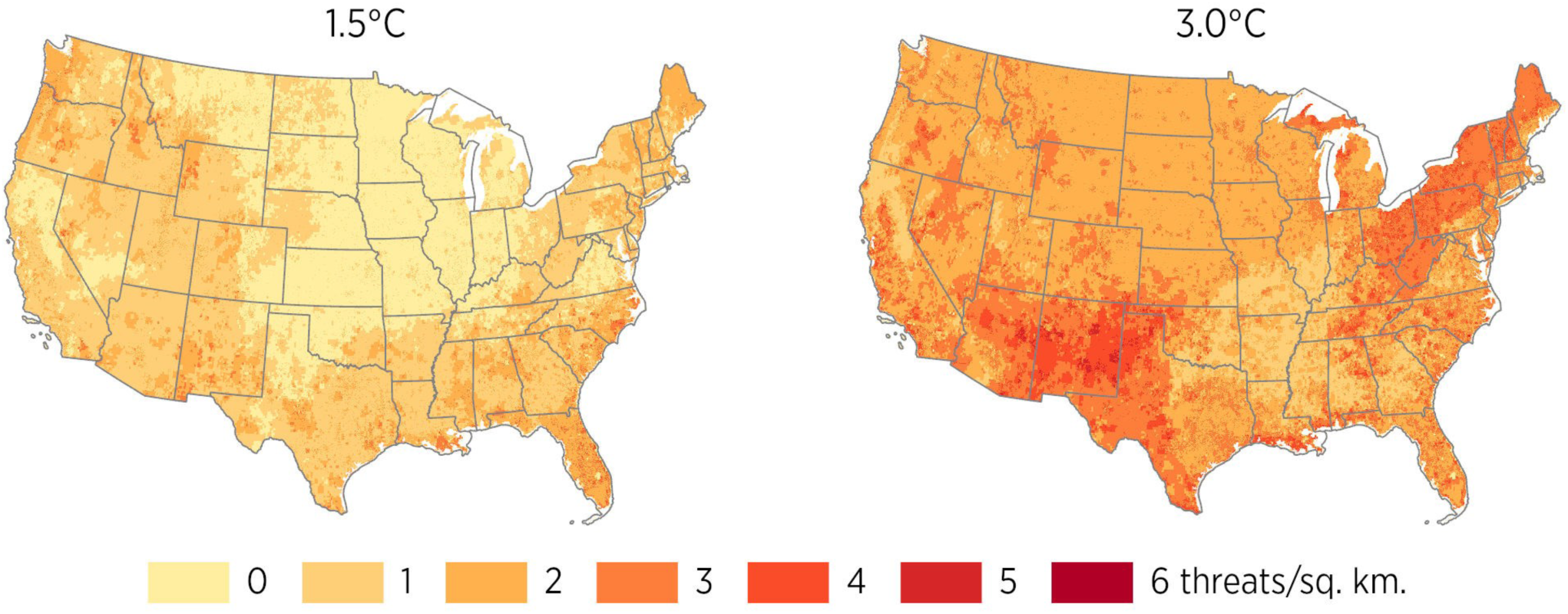
Coincident threats under future global change scenarios of 1.5°C and 3.0°C. For threats representing the proportion of each grid cell affected, we classified layers into binary outputs based on a threshold of >=25% of the cell and then summed these layers together to calculate the coincident number of threats.

Under this scenario, coincident threats were most concentrated in the coastal Southeast (Carolinas, Florida, and Louisiana), Pacific Northwest (western Oregon and Washington), and Intermountain West (Idaho, Wyoming, and New Mexico), and least concentrated in the Midwest and northern Great Plains (Fig 4). Coincident threats were relatively less concentrated in Northern California and Virginia across both scenarios (Fig 4).

### Birds: Exposure and Vulnerability

Spatial patterns of future bird species richness differed across seasons (S2 Fig), with richness projected to be more uniformly distributed in the breeding season and concentrated towards southern and coastal regions in the non-breeding season. During the breeding season, richness was greatest in mountainous regions of the West (i.e. the Sierra Nevadas, Cascades, and Rockies), the Upper Midwest, Northeast, and Southern Texas. During the non-breeding season, richness was greatest in low-lying and coastal regions, including the Southeastern Plains, Gulf Coast, California Coast, Central Valley, Willamette Valley, and Puget Trough (S2 Fig). Richness is projected to be greater in the non-breeding season than in the breeding season, and under 1.5°C than under 3.0°C. Looking across seasons and scenarios, the maximum projected richness was 235 non-breeding species/ km2 under the 1.5°C warming scenario.

Vulnerable species, those with high or moderate vulnerability scores based on range loss and gain [66], are projected to be coincident primarily along mountain ranges in the West. In the breeding season, vulnerability was highest in the Sierra Nevadas, Cascades, and Rockies under both scenarios, and also in the Northeast and Upper Midwest under 3.0°C. In the nonbreeding season, vulnerability was highest in the Pacific Coast Range, Sierra Nevadas, Cascades, Rockies, and South Florida under both scenarios (S3 Fig). Unlike overall species richness, vulnerability is projected to be greater in the breeding season than in the non-breeding season, and under 3.0°C than under 1.5°C. Maximum vulnerable species richness across seasons and scenarios was 58 breeding species/km2 under a 3.0°C warming scenario.

### Places at Risk

Risk, highlighting areas of high hazard (cumulative threats), exposure (bird richness), and vulnerability (vulnerable bird richness), was greater under 3.0°C compared to 1.5°C, and during the breeding season compared to the non-breeding season (Fig 5a,b). More vulnerable species and increased threats at 3.0°C translates to higher risk in both the breeding and nonbreeding seasons. In the breeding season, risk was greater across over 91% of the conterminous US under 3.0°C compared to 1.5°C, with more than a 100% increase across 70% of the US (Fig 5c). Under both scenarios, risk was greatest in mountainous regions of the West, but increased sharply in the Northeast and upper Midwest under 3.0°C (Fig 5). Risk decreased across less than 9% of the US under 3.0°C compared to 1.5°C in the breeding season, particularly in western Washington and Oregon, and in small patches across the Intermountain West, Coastal Plains, and Nebraska due to both lower hazard and fewer species present in these areas.

**Fig 5.**
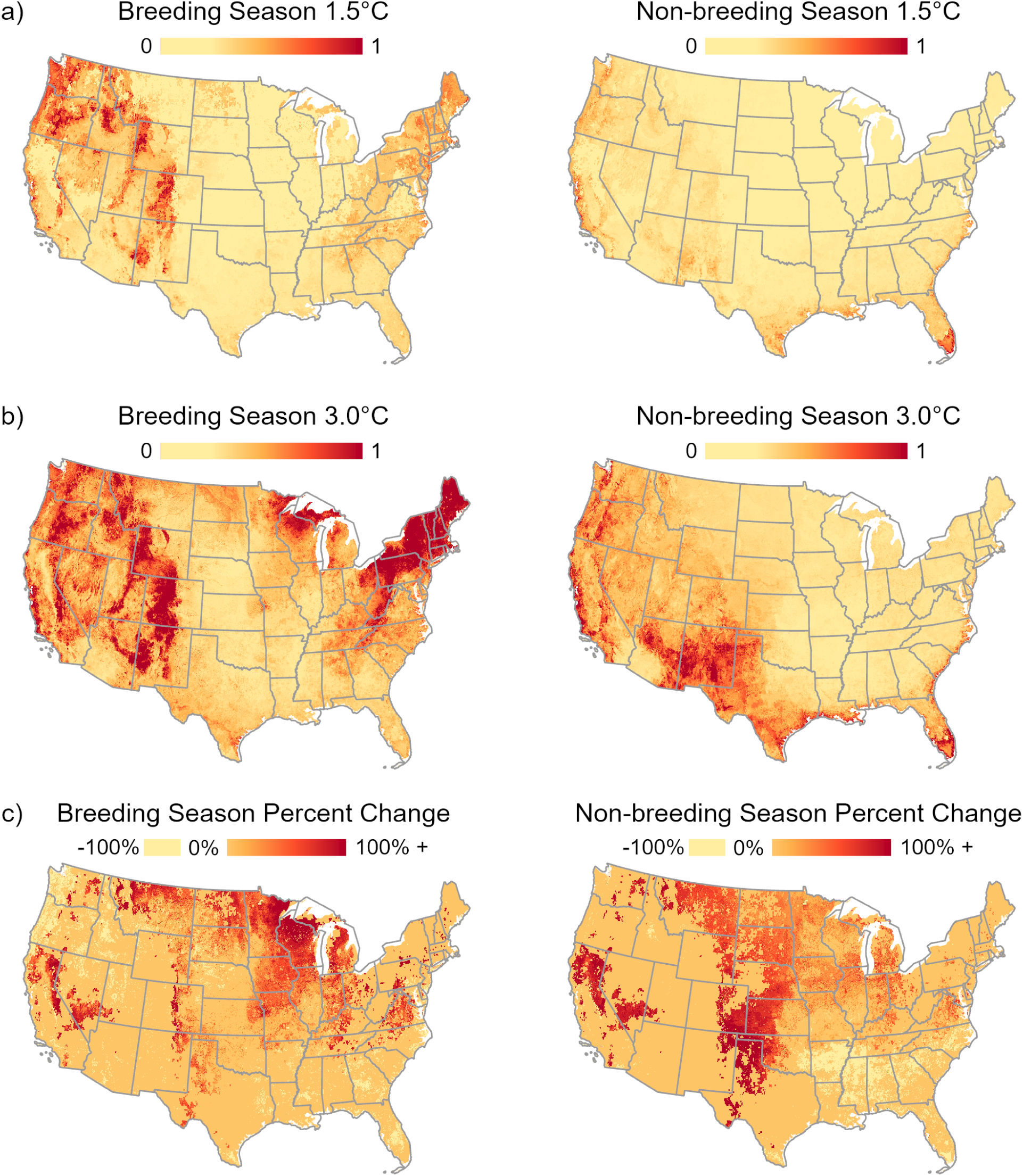
Risk to birds in the breeding and non-breeding season under future global change scenarios of 1.5°C and 3.0°C. Risk is calculated as the product of hazard (coincident threats), exposure (bird richness), and vulnerability (vulnerable bird richness) and then rescaled using min-max normalization. For each season, percent change between scenarios was calculated as the difference between risk under 3.0°C and 1.5°C divided by risk under 1.5°C.

In the non-breeding season, risk remained relatively high in the mountainous West, but was more pronounced farther south in New Mexico, Texas, the Gulf Coast, and Southeast Atlantic (Fig 5). As in the breeding season, risk was distributed fairly similarly across scenarios, but increased across 93% of the conterminous US under 3.0°C, including more than a 100% increase across 82% of the US. The Great Plains, Northern California, and Southern Nevada saw the largest increases in risk. Risk decreased across 7% of the US in the non-breeding sesaon in a few areas along the northern extent of the Coastal Plains and in the southern Interior Lowlands due to both lower hazard and fewer vulnerable species present in these areas.

### Impacts on Species

Extreme spring heat posed the greatest risk to birds in terms of number of species threatened, affecting 99% of species in both the breeding and non-breeding season under a 3.0°C warming scenario, including nearly 99% of vulnerable breeding species and 100% of vulnerable non-breeding species. Fire weather was the second greatest threat under 3.0°C, affecting just over 70% of species in the breeding season and nearly 50% of species in the non-breeding season, including 78% of vulnerable breeding species and 56% of vulnerable non-breeding species. Other extreme weather events were projected to have a substantial impact as well (Fig 2). Under 3.0°C, spring droughts were estimated to affect 20% of breeding and 26% of non-breeding species, including nearly a quarter of vulnerable non-breeding species, while heavy rains were estimated to affect 13% of breeding and 14% of non-breeding species, including approximately one-fifth of vulnerable species in both seasons. False springs posed the least threatening change among extreme weather events, affecting less than 1% of species across seasons; however, the few species affected were all vulnerable.

Extreme spring heat was also the greatest threat under 1.5° C, affecting 67% of breeding species and 82% of non-breeding species, including 73% of vulnerable breeding species and 97% of vulnerable non-breeding species. In general, however, impacts of extreme weather events were reduced under 1.5°C, with no species affected by fire weather or spring drought in either season, any only one species affected by false springs in the breeding season under this scenario. Heavy rain, however, still affected 9% of breeding and 10% of non-breeding species, including 15% of vulnerable breeding species and 20% of vulnerable non-breeding species.

Persistent threats generally had a smaller impact on birds, but urbanization and sea level rise both had disproportionate impacts on species relative to their extent (Fig 2). Although projected to cover 10% of the conterminous US under 3.0°C, urbanization was among the greatest threats to birds, affecting 44% and 61% of species in the breeding and non-breeding seasons. These impacts were greatly reduced under 1.5°C, with less than 3% of both area and species affected. Sea level rise had a smaller, but much more disproportionate, impact. Although it covered less than 1% of the conterminous US, sea level rise was projected to affect 4-5% and 9% of breeding and non-breeding species based on median estimates under both 1.5°C and 3.0°C. Considering high-end sea level rise estimates under 3.0°C, the number of species affected increased to 6% and 14% in the breeding and non-breeding seasons. Other persistent threats, including cropland expansion and lake level change, were projected to have minimal impacts.

### Coincident Threats

The majority of species analyzed were projected to be affected by at least one threat in each season and scenario, with up to four threats impacting a single species (Fig 3). Under 3.0°C, the vast majority of species were affected by multiple threats, with 97% of breeding species and 96% of non-breeding species experiencing two or more threats. The non-breeding season under 3.0°C had the most species affected by the greatest number of threats, with 57 species (12%) affected by four threats, while the breeding season under 1.5°C had the most species without any impacts (n=142, 32% of species analyzed). Reducing warming to 1.5°C, would result in most species projected to experience only a single threat (58% of breeding species, 64% of non-breeding species). Impacts on vulnerable species were also greater under a 3.0°C scenario, with at least 94% of vulnerable species facing multiple threats, and 12% of vulnerable species facing four threats. However, if we stabilize warming at 1.5°C globally through emissions reduction, at least 63% of vulnerable species would face only a single threat across seasons, and up to 24% of vulnerable species would not face any threats in the breeding season.

Some habitat groups were more likely to face multiple threats than others (Fig 6). In both seasons under 3.0°C, the coastal group had the most species facing four coincident threats. The subtropical forests and marshland groups also had a high proportion of species facing four threats in both seasons, along with eastern forests in the breeding season, and aridlands and western forests in the non-breeding season. Under the 3.0° C scenario, groups that had more than 50% of species affected by three or more coincident threats in the breeding season included coastal (69%), eastern forests (60%), subtropical forests (93%), aridlands (60%), and urban/suburban (88%). In the nonbreeding season, these groups included coastal (78%), aridlands (81%), subtropical forests (96%), boreal forests (54%), and urban/suburban (75%). Across groups, the number of coincident threats was greatly reduced under 1.5°C (S4 Fig). At this lower climate change scenario, the majority of groups had 50% or more of species facing 0-1 threats (S4 Fig).

**Fig 6.**
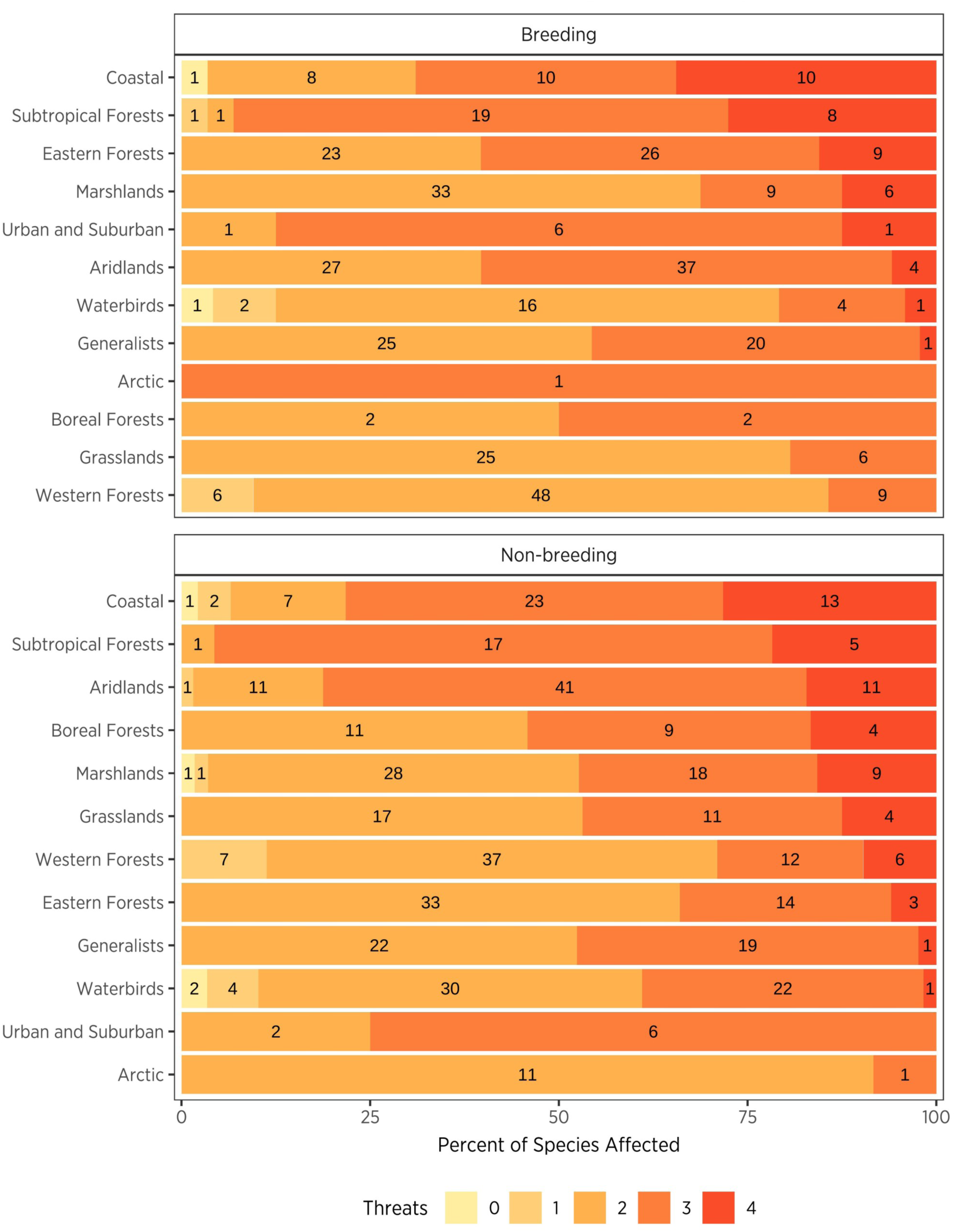
Proportion and number of (a) breeding and (b) non-breeding species per bird habitat group affected by coincident number of threats under a 3.0°C future global change scenario. Groups are listed in descending order by the total number of species that face four threats.s

## DISCUSSION

### Patterns of Climate-Related Threats

Our results indicate that with unmitigated climate change, over 88% of the conterminous US will be affected by multiple coincident threats, and the additive nature of these threats compounds the stress climate change already has on biodiversity [3,10]. Climate-related threats are more widespread and intense under a 3.0°C scenario, and some locations might experience up to six of the nine threats included in this analysis. Indeed, under the higher 3.0°C scenario, at least 96% of species are projected to experience multiple coincident threats across seasons, and more than 50% of species within the coastal, subtropical forests, and urban/suburban habitat groups will face 3-4 coincident threats in both seasons. 98% of the conterminous 48 states could be affected by one or more climate-related threats, and 97% of species could be affected by two or more climate-related threats. Furthermore, at least 70% of the conterminous US faces a greater than 100% increase in risk between scenarios, indicating that the majority of the lower 48 states will likely see unprecedented risk for birds and the places they need. However, emissions mitigation to reduce warming to 1.5°C would reduce risk across over 90% of the conterminous US in both seasons. These findings support the IPCC recommended target of a 1.5°C-2.0°C global mean temperature rise to minimize climate change effects [24]. Given that we have already seen a near 1.0°C increase globally over pre-industrial levels [24] and potential warming likely will exceed this (4-5+ °C) by the end of century [82,83], we have limited time to act.

Comparing seasons, climate change risk for birds was higher in the breeding season compared to the non-breeding season, despite higher exposure (i.e. species richness) in the nonbreeding season. This pattern was driven by higher species vulnerability in the breeding season, related to high rates of species range loss and northward range shifts in the conterminous US [66]. We observe heightened risk in the Northeast and Upper Midwest as more bird species shift into this area at 3.0°C, increasing both exposure and vulnerability. Changes in climate during the breeding season can influence species persistence by regulating breeding productivity [84–87], and could potentially lead to reduced breeding success, population declines, and local extinction [88]. Birds with range shifts, such as those moving into the upper Midwest and Northeast, will need to cope with these additional threats as they seek newly suitable climatic conditions, or face altered climate patterns in areas where they remain, all the while facing new, coincident threats.

In the non-breeding season, risk is highest along the Gulf, Southwest, and Southeast Atlantic US at 3.0°C due to high exposure and hazard, as these regions see high species richness and multiple threats. These areas also see substantial gains in species shifting and expanding their ranges [66]. Mortality is already high for wintering and resident species in these areas [89], and additional pressure from climate change and multiple compounding threats could curtail species range shifts and reduce anticipated species richness in these areas. Risk was also notably high in the mountainous regions of the West across seasons and scenarios, areas that have both high species richness and species vulnerability. Mountains may become critical climate refugia [90] and corridors for species shifting ranges in response to climate [91]. However, mountainous regions are particularly sensitive to climate change, as climate isoclines shift upslope and can eventually disappear if the rate of warming is too high (La Sorte & Jetz, 2010; Laurance et al., 2011; Sekercioglu et al., 2008). Finally, no area in the conterminous US is devoid of risk, and areas that see less risk comparatively (e.g., the central Midwest) exhibit the greatest increase in risk between 1.5°C and 3.0°C, highlighting that these areas will become more at risk with increased warming.

### Intermittent vs. Persistent Threats

Across the threats analyzed here, short-term intermittent threats (i.e. extreme weather events) had the greatest influence on risk and the widest spatial coverage. Although these threats are historically uncommon, both their magnitude and frequency are anticipated to increase with climate change [43,44,94]. Indeed, extreme spring heat occurring every two years or more often are set to affect nearly the entire conterminous US at the 3.0°C scenario, overlapping with 99% or more of species. Historically, these events were only seen every 20 years or more [43]. The wide-ranging extent and frequent timing of extreme spring heat in the early breeding season can increase heat stress on birds during a critical period, reducing population growth [27,95] and increasing bird mortality [28,96]. Extreme weather in one season can also have cascading effects on other parts of the year, and can have a delayed effect on habitat quality and populations [32,97,98].

In addition, increased fire weather, a measure of drought indicative of wildfire potential, is projected to affect two-thirds of the country at the 3.0°C scenario. This pattern indicates that the majority of the conterminous US will have one-quarter of the year or more in weather conditions that are ideal for wildfires and drought, a more than 233% increase from historical levels [44]. Fire weather will also affect more than half of species under 3.0°C, and is skewed to areas with high species vulnerability, affecting 78% of breeding and 56% of non-breeding vulnerable species. The relationship between fire and birds is complex, and natural wildfire regimes have been altered to the detriment of fire-adapted systems due to the legacy of fire suppression; at the same time, studies find pyrodiversity (i.e. a diversity of fire severities) might benefit bird diversity (Tingley, Ruiz-Gutiérrez, Wilkerson, Howell, & Siegel, 2016; Smucker, Hutto, & Steele, 2005). With climate change, however, there is evidence for a regime shift from the natural system of heterogeneous mixed-severity fires being replaced with large and homogeneous high-severity fires [100,101]. These larger, more severe fires can have a negative effect on bird abundance [102], and may lead to habitat loss and delayed habitat regeneration in the long-term [103,104].

Heavy rains and spring droughts are also expected to increase, with nearly one-quarter of the country and over 20% and 13% of breeding species affected by each threat, respectively. These threats are mostly distinct in their spatial coverage, however, with spring droughts concentrated in the Southwest and heavy rains in the East and Pacific Northwest. Both of these types of extremes in precipitation are known to cause mortality events in birds [27], as well as reduced reproductive success [105]. For heavy rainfall, this is related to decreased parental nest visitation, diminish parental survival, and subsequently reduces recruitment [106], as well as damage and inundation of nests leading to mortality [27,34–38]. Mortality and reduced reproduction from droughts are linked with nest abandonment and skipped breeding [31], which can be exacerbated by heat stress [45,96]. Changes in precipitation can also have a cascading effect through food webs, further stressing species and ecosystems [107]. The detrimental effects of these short-term intermittent threats paired with their extensive spatial coverage, increased frequency, and coincident nature at the higher climate change scenario will be challenging for birds.

Although intermittent threats covered large spatial areas, the persistent and catastrophic threats of urbanization and sea level rise will disproportionately affect bird species relative to their spatial coverage. Urbanization is one of the greatest threats to birds. Although only covering 10% of the conterminous US, we see nearly half of breeding species and more than half of nonbreeding species affected. This finding includes many vulnerable species, including one-third in the breeding season and nearly half in the non-breeding season. The reason for this disproportionate effect could be that areas that are attractive to humans for development are often areas of high suitability and potential richness for birds [108,109]. This indicates that birds and humans favor similar environments at some level, but high human density and land use suppresses bird abundance and species richness [59,109–111]. With unmitigated climate change, we anticipate increased urbanization, modification, and fragmentation of habitat from human land use [55,56]. Here, we find that urbanization will coincide with areas suitable for many bird species, including areas that will become newly suitable, likely limiting the ability of species to shift their ranges and find appropriate habitat not already altered by humans [108]. Unless there are measures to stabilize climate change to 1.5°C and reduce this threat, urbanization will exacerbate a stressed system that has already seen declines in species richness and abundance associated with anthropogenic land use [58–61].

In addition, sea level rise will affect 3-13% more species than the proportion of area affected. Coastal habitats support large numbers of bird species in concentrated areas along shorelines, beaches, and tidal areas, and sea level rise is anticipated to increase erosion, shoreline retreat, and loss of coastal habitat [112–114]. In addition, coastal systems are increasingly faced with encroachment from coastal development and increased housing density, population growth, and disturbance [115]. The combination of habitat loss from inundation and erosion due to sea level rise, and coastal armoring to counteract these impacts, can lead to loss of roosting sites, lower species abundance and richness, and higher mortality and nest failures [25,112]. In addition, land development can limit areas that coastal habitat and marshes can migrate into as sea levels rise [48].

### Greater Risk for Certain Groups

Of note, the habitat groups with the greatest number of species facing coincident threats (subtropical forests and coastal, which were classified as intermediate vulnerability; and urban/ suburban and marshlands, which were classified as low vulnerability) were not amongst the most vulnerable groups based on previous assessment from projected range shifts. Species in these habitat groups may see relatively stable ranges (with more areas remaining or becoming suitable than losing suitability), but when faced with multiple climate change associated threats, they may need additional conservation efforts to ensure persistence. As coincident threats can act in synergy, potentially as feedback, accelerating each other [12], species and groups facing multiple threats may be at higher risk to climate change impacts. For forest species, this is true of land use change and warming and drought working in combination to amplify and expedite bird population declines [12]. Additionally, wildfires may also become more frequent with increased human populations and encroachment on wild areas [116], further altering natural fire regimes and adding potential sources of ignition, a source of fire risk that we were unable to project in our analysis. For coastal systems, increased pressures from both urbanization, causing habitat loss inland [115,117], and climate-driven sea level rise, causing habitat loss from the oceans, may create a squeezing effect that accelerates decline in these systems. These examples highlight how researchers and managers can use threats analyses such as this one to more completely assess vulnerability and risk to species and habitats under climate change [11].

### Caveats

Despite most of our species’ distributions covering a much wider extent in North America, we decided to focus on a US-based risk assessment due to limitations in the availability and spatial coverage of our threats data. We also understand that multiple threats not addressed here could also affect birds under climate change scenarios, including increased exposure to pollution [e.g., pesticides;, 118], heavy metal depositing; [119], expansion of invasive species [120], increased predation pressure [e.g., snakes;, 88], exposure to novel or intensifying diseases [e.g., West Nile virus, tick-borne illnesses, Bird Flu;, 121], increasing ocean temperatures [122], food chain collapse [123], or altered biotic interactions [124,125], to name a selection of potential threats.

Spatial resolution may also have exaggerated our estimated extent and percent of range affected by threats. Persistent threats were available at a 1-km resolution or finer, while extreme weather data were only available at a 12-km resolution. Our extreme weather events covered the largest area, and the coarse inputs may confound estimates of area affected. This effect is likely exacerbated in topographically complex locations. However, climate conditions do operate at broad spatial scales, and a 12-km resolution likely captures enough spatial information to guide on the nature of the extremes locally. Additionally, we did not resolve potentially overlapping and conflicting projections (e.g., flooding by sea level rise and urbanization) for the same cell. This decision likely has a minimal effect on the risk analysis, but may lead to overestimates where only one threat would be able to occur (e.g., cropland expansion and urbanization could be mutually exclusive). Lastly, we based our threat impacts on species range area, which does not account for variable patterns of species abundance throughout a range, potentially neglecting areas that may be more important for a species’ conservation due to higher relative abundance.

## CONCLUSION

Our results highlight that bird species in the US are at risk from both range shifts and persistent and intermittent threats related to climate change; over 88% of the conterminous US and at least 96% of species will be affected by multiple coincident threats that are traditionally not considered in species distribution models. The added layer of these, and other, climate-related threats amplifies the risk species already face in ongoing climate change. Regardless of season, climate-related threats are more widespread and intense at the 3.0°C scenario compared to the 1.5°C scenario, but climate change mitigation would reduce risk to birds from climate change-related threats across over 90% of the conterminous US. Our maps of risk and lists of species and groups most affected by these threats will inform conservation planning and climate change adaptation for birds and the places they need.

## ACKNOWLEDGMENTS

This work was funded by the John D. and Catherine T. MacArthur Foundation, grant G-1511-150388 and by U.S. Fish and Wildlife Service funding 140F0318P0263. We also received support from Amazon Web Services’ (AWS) Promotional Credit Program (https://aws.amazon.com/awscredits/) and Domino Data Labs Programs for Non-profits and Education (https://www.dominodatalab.com/domino-for-good/) for data science platform use and cloud processing. We thank Andrew Allstadt and the SILVIS lab at the University of Wisconsin-Madison for providing support and data processing. The findings and conclusions in this article are those of the author(s) and do not necessarily represent the views of the U.S. Fish and Wildlife Service. To the extent possible, this research was completed in compliance with the Guidelines to the Use of Wild Birds in Research; however, the authors were not directly involved in the collection of the datasets used.

## SUPPORTING INFORMATION

**S1-4**. Supplementary Figures.

