## Supplementary material for "Risk to North American Birds from Climate Change-Related Threats": S1-4 Supplementary Figures

### Supporting Figures S1-S4

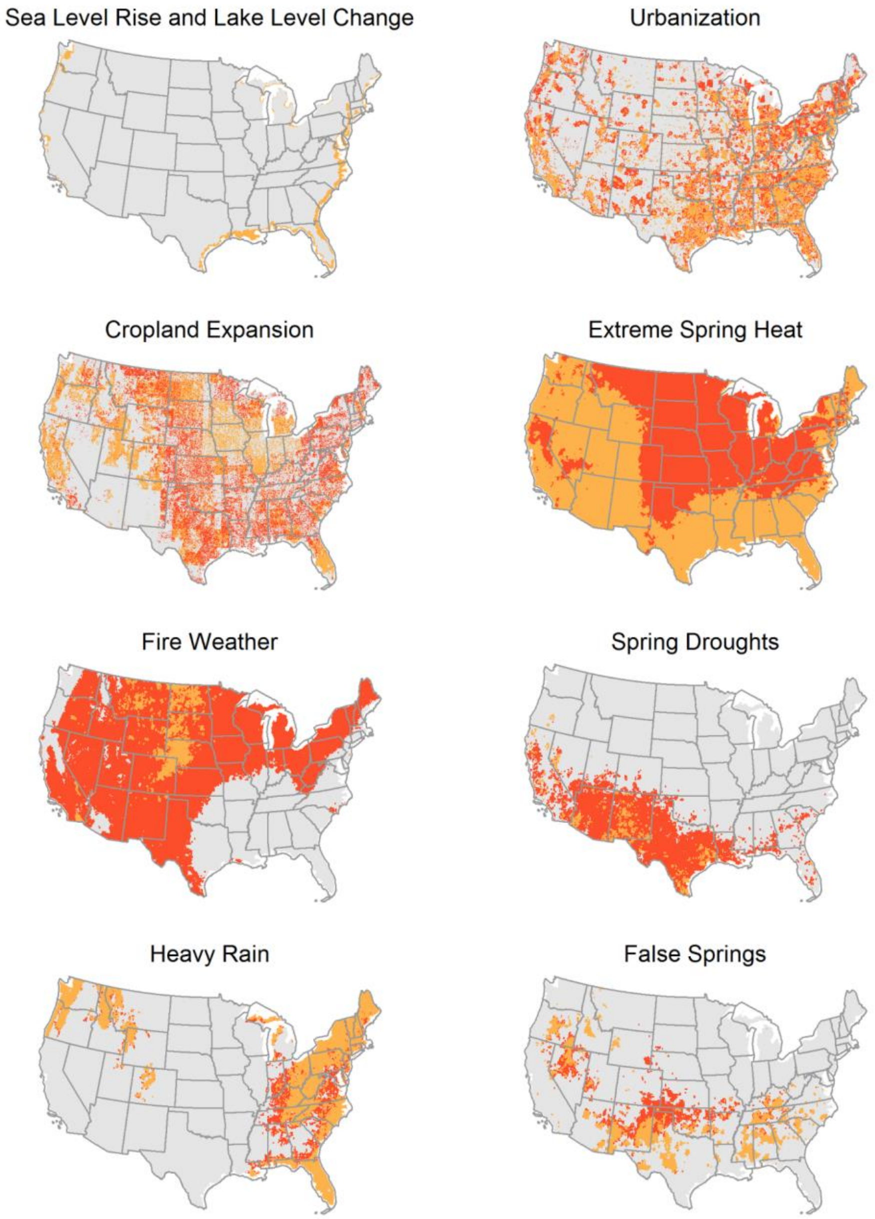


**S1 Fig. Threats under future global change scenarios of 1.5°C and 3.0°C.** For threats representing the proportion of each grid cell affected (i.e. sea level rise, lake level change, and urbanization), we classified layers into binary outputs based on a 25% threshold. In each map, threats are shown in orange under 1.5°C and red under 3.0°C. Areas in red can be interpreted as the additional threat under 3.0°C, as nearly all threats covered more area under this scenario, except cropland expansion, where the pattern is reversed, and orange areas represent the additional threat under 1.5°C.

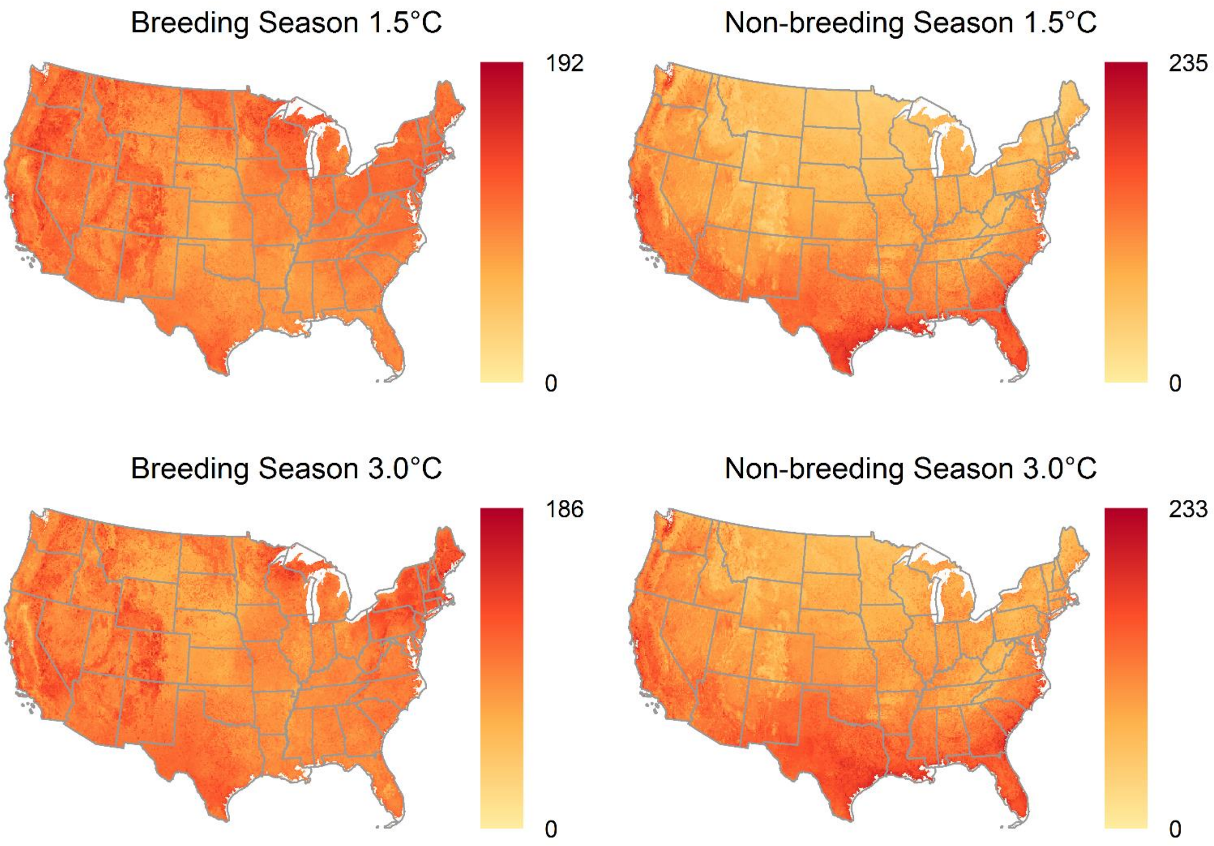


**S2 Fig. Bird richness in the breeding and non-breeding season under future global change scenarios of 1.5°C and 3.0°C.** Bird richness is calculated as the sum of modeled future ranges for species with at least 10% of their range in the conterminous US.

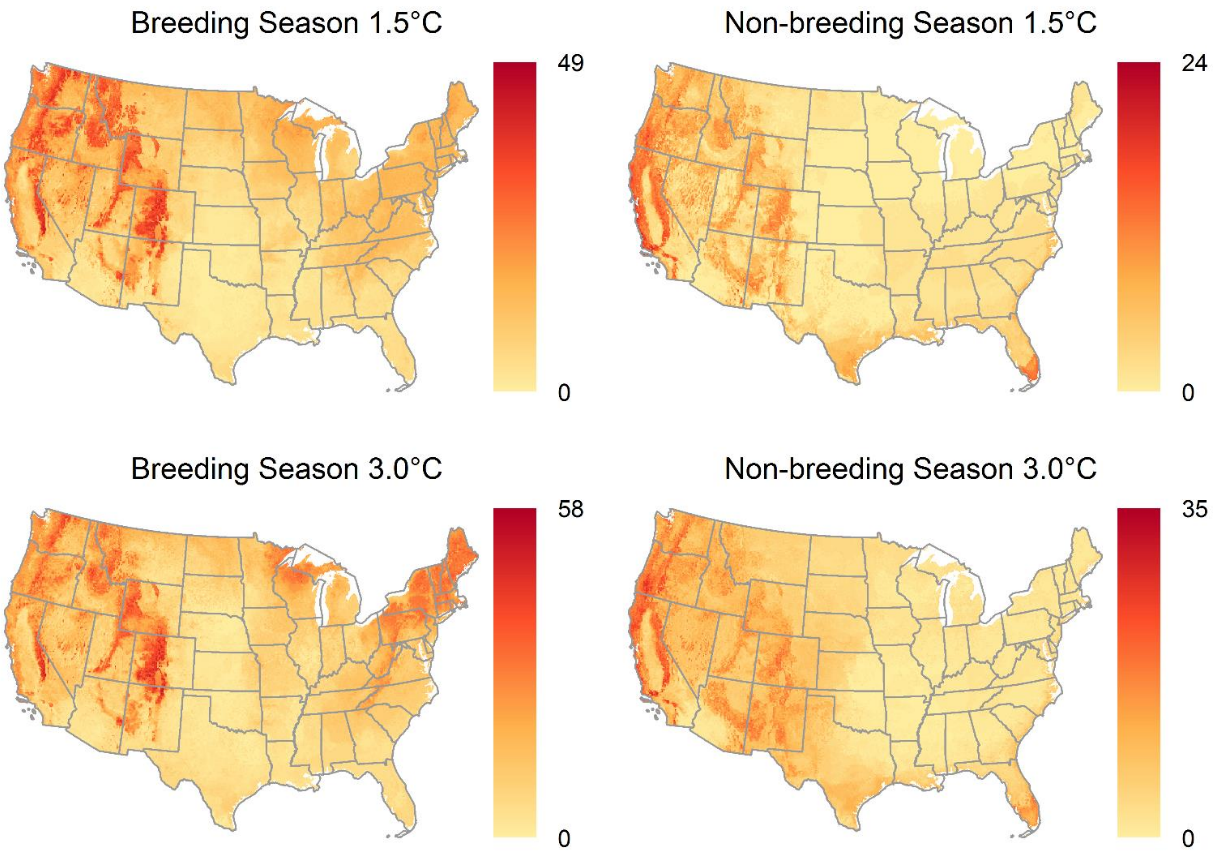


**S3 Fig. Bird vulnerability in the breeding and non-breeding season under future global change scenarios of 1.5°C and 3.0°C.** Bird vulnerability is calculated as the sum of modeled future ranges for species that were classified as moderately or highly vulnerable to climate change and had at least 10% of their range in the conterminous US.

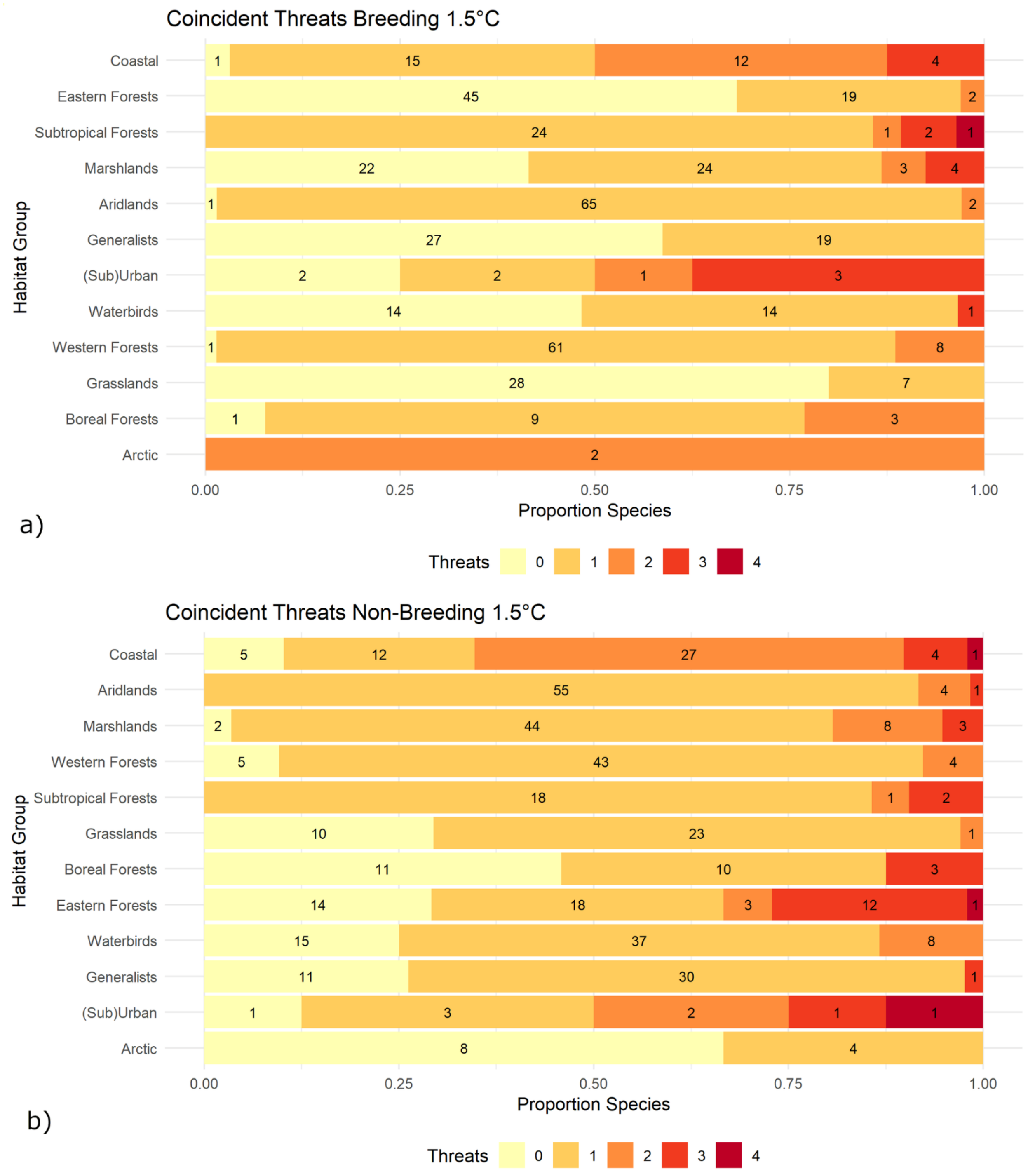


**S4 Fig. Proportion and number of (a) breeding and (b) non-breeding species per bird habitat group affected by coincident number of threats under a 1.5°C future global change scenario.** Groups are listed in the same order as Fig 6 in the main document.
